# Clinical performance of SARS-CoV-2 IgG antibody tests and potential protective immunity

**DOI:** 10.1101/2020.05.08.085506

**Authors:** Niko Kohmer, Sandra Westhaus, Cornelia Rühl, Sandra Ciesek, Holger F. Rabenau

**Affiliations:** Institute for Medical Virology, University Hospital, Goethe University Frankfurt am Main, Frankfurt, Germany; German Centre for Infection Research, External partner site Frankfurt, Germany

## Abstract

As the current SARS-CoV-2 pandemic continues, serological assays are urgently needed for rapid diagnosis, contact tracing and for epidemiological studies. So far, there is little data on how commercially available tests perform with real patient samples and if detected IgG antibodies provide protective immunity. Focusing on IgG antibodies, we demonstrate the performance of two ELISA assays (Euroimmun SARS-CoV-2 IgG & Vircell COVID-19 ELISA IgG) in comparison to one lateral flow assay ((LFA) FaStep COVID-19 IgG/IgM Rapid Test Device) and two in-house developed assays (immunofluorescence assay (IFA) and plaque reduction neutralization test (PRNT)). We tested follow up serum/plasma samples of individuals PCR-diagnosed with COVID-19. Most of the SARS-CoV-2 samples were from individuals with moderate to severe clinical course, who required an in-patient hospital stay.

For all examined assays, the sensitivity ranged from 58.8 to 76.5% for the early phase of infection (days 5-9) and from 93.8 to 100% for the later period (days 10-18) after PCR-diagnosed with COVID-19. With exception of one sample, all positive tested samples in the analysed cohort, using the commercially available assays examined (including the in-house developed IFA), demonstrated neutralizing (protective) properties in the PRNT, indicating a potential protective immunity to SARS-CoV-2. Regarding specificity, there was evidence that samples of endemic coronavirus (HCoV-OC43, HCoV-229E) and Epstein Barr virus (EBV) infected individuals cross-reacted in the ELISA assays and IFA, in one case generating a false positive result (may giving a false sense of security). This need to be further investigated.

## Background

SARS-CoV-2 is a new Coronavirus, belonging to the group of betacoronaviruses, which emerged in December 2019 in Wuhan, China. It is the causative agent of an acute respiratory disease known as coronavirus disease 2019 (COVID-19). The spectrum of clinical signs can be very broad and asymptomatic infections are reported. The virus has rapidly spread globally. On 11 March 2020 the World Health Organization (WHO) declared COVID-19 as a pandemic. Nucleic acid amplification testing (NAT) is the method of choice in the early phase of infection (1). However, to acquire knowledge about the seroprevalence of SARS-CoV-2 and to test for (potential) individual immunity, there is an increasing demand in the detection of antibodies – especially of IgG antibodies. Convalescent plasma may be used for therapeutic or prophylactic approaches as vaccines and other drugs are under development (2). For all these purposes, sensitive and especially highly specific antibody assays are needed. The spike (S) protein of SARS-CoV-2 has shown to be highly immunogenic and is the main target for neutralizing antibodies (3). Currently there are many S protein based commercially or in-house developed assays available, but there is limited data on how these tests perform with clinical samples and if the detected IgG antibodies provide protective immunity. This study aims to provide a quick overview on some of these assays (two commercial available ELISA, an LFA, an IFA and a PRNT, focusing on the detection and neutralization capacity of IgG antibodies in follow up serum or plasma samples of individuals with PCR-diagnosed infections with SARS-CoV-2. To assess potential cross-reactivity, we examined defined follow-up samples of individuals infected with endemic coronaviruses and other infectious diseases.

## Materials and methods

### Serum and plasma samples

We collected follow up serum or plasma samples (in the following simply stated as samples) from individuals with PCR-diagnosed infections with SARS-CoV-2 (n=33) at different time points (table 1). Most of these individuals had a moderate to severe clinical course and required an in-patient hospital stay at the intensive care unit. Additionally, follow up samples of recent PCR-diagnosed infections with SARS-CoV (3 patients from the 2003 outbreak), HCoV-OC43 (n=4), HCoV-HKU1 (n=1), HCoV-NL63 (n=2), HCoV-229E (n=4) and recent serological/PCR-diagnosed infections with acute EBV (n=4, three serologically EBV-VCA-IgM positive and one PCR- and serologically EBV-VCA-IgM positive) and acute CMV (n=3) (all serologically IgM and PCR-positive) were collected. The samples of individuals infected with endemic human coronavirus, CMV and EBV were used to assess potential cross reactivity and the risk of potential false positive results.

**TABLE 1.** Sensitivity and specificity of the examined SARS-CoV-2 IgG assays from days 5-9 and days 10-18.

| Company | Days after confirmed SARS-CoV-2 PCR |  |  |  |
| --- | --- | --- | --- | --- |
|  | 5-9 (days) | 10-18 (days) |  |  |
|  | sensitivity (%) |  | specificity (%) | specificity (%) incl. SARS-CoV (2003)* |
| Euroimmun (ELISA) | 58.8 (10/17) | 93.8 (15/16) | 95.7 (22**/23) | 96.2 (25/26) |
| Vircell (ELISA) | 70.6 (12/17) | 100 (16/16) | 95.2 (20/21) | 83.3 (20/24) |
| IFA (in-house) | 76.5 (13/17) | 100 (16/16) | 100 (19/19)*** | 86.4 (19/22) |
| Assure Tech (Rapid test) | 62.5 (10/16) | 93.8 (15/16) | 100 (13/13) | - |
| PRNT (in-house) | 76.5 (13/17) | 100 (16/16) | - | - |

### Lateral flow assay

The FasStep (COVID-19 IgG/IgM) rapid test cassettes (COV-W32M, Assure Tech (Hangzhou) Co., Ltd, China) were used according to the manufacturer’s recommendation. We have no details on the used antigen component. 10 µl serum and two drops of sample buffer were applied to the sample well. Test results were visually evaluated after 10 minutes.

### ELISA

The CE certified versions of the Euroimmun SARS-CoV-2 IgG ELISA (Euroimmun, Lübeck, Germany) and Vircell COVID-19 ELISA IgG (Vircell Spain S.L.U., Granada, Spain) were used, in an identical manner, according to the manufacturer’s recommendation. Both ELISAS use SARS-CoV-2 recombinant antigen from spike glycoprotein (S protein) and the Vircell ELISA additionally Nucleocapsid (N protein). Samples were diluted 1:101 or 1:20, respectively, in sample buffer and incubated at 37° for 60 min in a 96-well microtiter plate followed by each protocols washing and incubation cycles, including controls and required reagents. Optical density (OD) was measured for both assays at 450 nm using a Virclia microplate reader (Vircell Spain S.L.U., Granada, Spain). The signal-to-cut-off ratio was calculated and values expressed according to each manufacturer’s protocol.

### Immunofluorescence assay (IFA)

For an immunofluorescence assay Vero cells (african green monkey, ATCC CCL-81 (American Type Culture Collection, Manassas, Virginia, USA)) were infected with SARS-CoV-2 and harvested two days post infection. Briefly, cells were trypsinized and washed once with PBS before transferred onto a 10-well diagnostic microscope slides. After drying, cells were fixated with 100% ethanol for 10 minutes. Patient samples were diluted in sample buffer (Euroimmun AG, Lübeck, Germany) in a dilution of 1:50 and 30 µl applied per well. The slides were incubated at 37°C for 1 hour and washed three times with phosphate-buffered saline (PBS)-Tween (0.1%) for 5 minutes. 25 µl of goat-anti human fluorescein-labeled IgG conjugate was used as secondary antibody. The slides then were incubated for 30 minutes and washed three times with PBS-Tween for 5 minutes. The microscopic analysis was performed by 200-fold magnification using a Leica DMLS fluorescence microscope (Leica Mikrosysteme Vertrieb GmbH, Wetzlar Germany).

### Plaque reduction neutralization test (PRNT)

To test for neutralizing capacity of SARS-CoV-2 specific antibodies, Caco-2 cells (human colon carcinoma cells, ATCC DSMZ ACC-169 (American Type Culture Collection, Manassas, Virginia, USA)) were seeded on a 96-well plate 3-5 days prior infection. 2-fold dilutions of the test sera beginning with a 1:10 dilution (1:10; 1:20; 1:40; 1:80; 1:160; 1:320; 1:640 and 1:1280) were made in culture medium (Minimum essential medium, MEM; Gibco, Dublin, Ireland) before mixed 1:1 with 100 TCID50 (Tissue culture infectious dosis 50) of reference virus (SARS-CoV-2 FFM1 isolate). FFM1 was isolated from a patient at University Hospital Frankfurt who was tested positive for SARS-CoV-2 by PCR. Virus-serum mixture was incubated for one hour at 37°C and transferred onto the cell monolayer. Virus related cytopathic effects (CPE) were determined microscopically 48 to 72 hours post infection. To determine a potential neutralizing ability of patient serum, CPE at a sample dilution of 1:10 is defined as non-protective while a CPE at a dilution of >1:20, is defined as protective.

## Results

In the early phase of infection, from days 5-9 after PCR-confirmed infection with SARS-CoV-2, the in-house developed IFA and PRNT showed a sensitivity of 76.5% (13/17), the Vircell ELISA a sensitivity of 70.6% (12/17), the Assure Tech Rapid Test a sensitivity of 62.5% (10/16) and the Euroimmun ELISA a sensitivity of 58.8% (10/17). For the later period from days 10-18, the Euroimmun ELISA and Assure Tech Rapid Test showed a sensitivity 93.8% (15/16), the Vircell ELISA, IFA and PRNT of 100% (16/16) - (TABLE 1). For selected samples (SARS-CoV samples from the 2003 outbreak excluded, TABLE S2), the Euroimmun ELISA showed a specificity of 95.7%, generating a borderline result for the HCoV-OC43 sample, the Vircell ELISA of 95.2%, generating a positive result for HCoV-229E sample and the in-house developed IFA of 100% (an unspecific result for one EBV sample was excluded). Including the three SARS-CoV samples from the 2003 outbreak, the Euroimmun ELISA showed a specificity of 96.2% (not generating any cross-reactive results for the SARS-CoV samples), the IFA of 86.4% and the Vircell ELISA of 83.3% (both assays generating positive results for all three SARS-CoV samples). The Assure Tech Rapid Test did not generate any false positive results for the tested samples. None of the other tested samples cross-reacted in terms of generating borderline or false positive results.

The signal-to-cut-off (S/CO) ratios of the Euroimmun and Vircell ELISA and the corresponding PRNT titers for the tested samples are shown in FIG 1. In samples 3, 10 and 11, none of the examined assays (including the IFA and Assure Tech Rapid Test), detected SARS-CoV-2 antibodies. In sample 1, only the Vircell ELISA, in sample 4 and 19 only the Vircell ELISA and PRNT (including the IFA) detected antibodies. In samples 12 and 16, only the PRNT (and IFA) detected antibodies (in sample 16 with a titer <1:10). With exception of sample 1, all with the ELISA positive tested samples were also positive tested with the IFA. In the detection of antibodies, the IFA performed like the PRNT on all examined samples. All with the commercially available assays positive tested samples (except of sample 1) showed neutralizing properties in the PRNT (titer >1:20), indicating a potential protective immunity.

**FIG 1.**
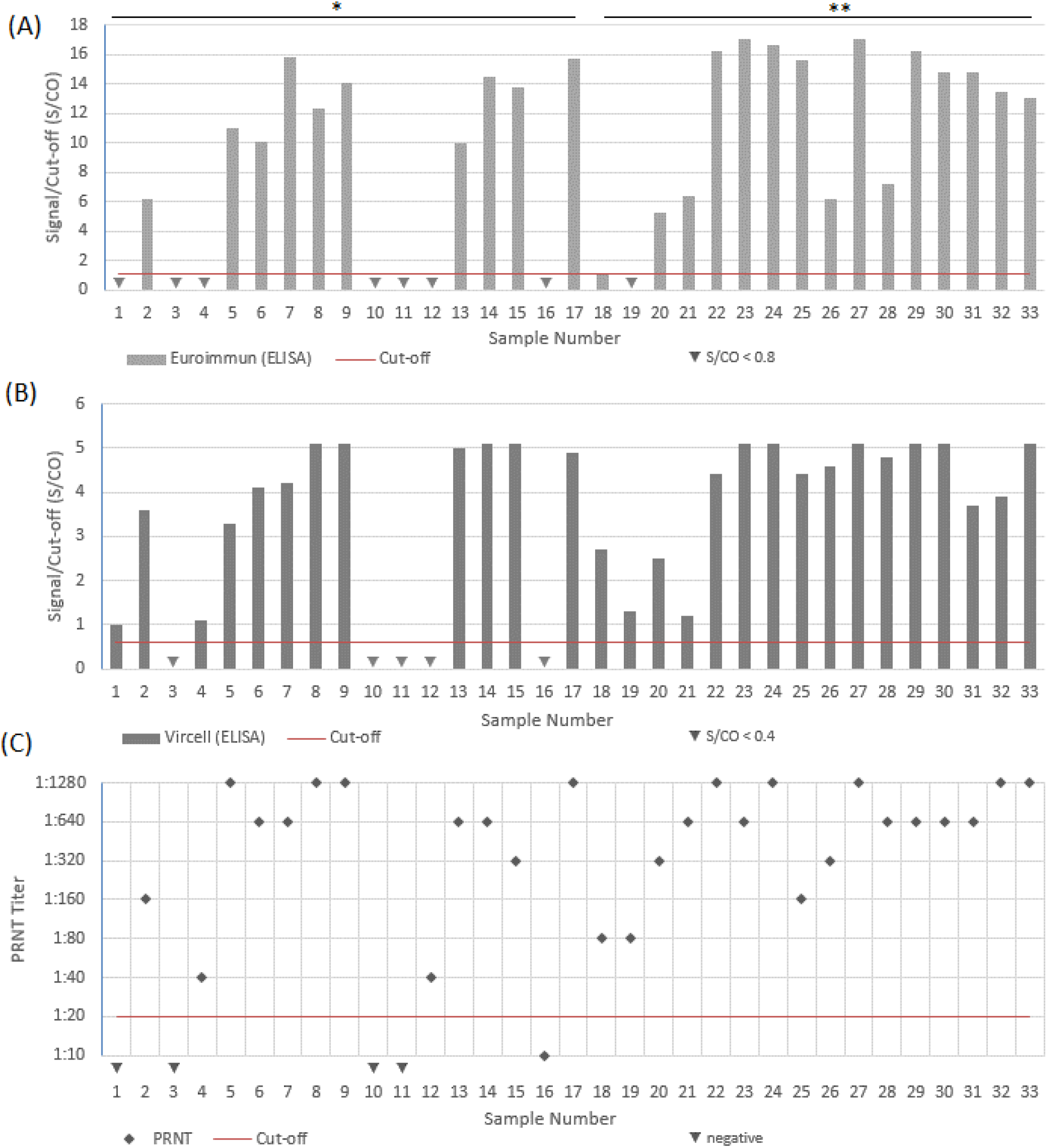
Results of the for sensitivity tested samples in the ELISA assays and PRNT; (A) Euroimmun ELISA Signal/Cut-off (S/CO) ratio of tested samples; (B) Vircell ELISA Signal/Cut-off (S/CO) ratio for tested samples; (C) PRNT Titer for tested samples. *Days 5-9 /**Days 10-18 after confirmed SARS-CoV-2 PCR.

## Discussion

In terms of sensitivity, our data are consistent with previously published data. In a study from Liu et al., using an rS-based ELISA assay, the group found SARS-CoV-2 IgG antibodies in less than 60% of the samples from days 6-10 after disease onset. The sensitivity increased to >90% in samples from days 16-20 (4). In a study from Wölfel et al., using an in-house developed IFA, the group found seroconversion in all examined follow-up serum samples of COVID-19 patients by day 14 after onset of symptoms. The samples were further analyzed via PRNT, all showed neutralization activity against SARS-CoV-2 (5).

An important finding of our study is, that (with exception of sample 1) all detected SARS-CoV-2 IgG antibodies in the analyzed cohort, using the commercially available assays examined, demonstrated neutralizing (protective) properties in the PRNT. The screening for SARS-CoV-2 IgG antibodies [especially for potential protective IgG antibodies against the S protein (6)] using ELISA or lateral flow assays is more convenient and practicable than using the hands on- and time-intensive IFA or PRNT, which can only be performed by experienced personnel, and the PRNT, only in a BSL-3 laboratory. ELISA based assays can be automated and used for larger sample sizes. Lateral flow assays can be used by less experienced personnel in a point-of-care setting, generating results in short time. Some samples, however, were only detected with the IFA and PRNT as gold standard. The titer needed for potential protective immunity is not yet (officially) defined. In one study, it is reported, that a individual cleared SARS-CoV-2 without developing antibodies up to 46 days after illness (7). The mechanism of immunity, especially of protective immunity (if applicable) and how long it will last, need to be further investigated. Besides humoral mediated immunity, there is evidence that T-cell mediated immunity plays a role (8). Most of the SARS-CoV-2 samples analysed in this study were from individuals with moderate to severe clinical course, who required an in-patient hospital stay. We have also tested follow-up samples of individuals PCR-diagnosed with COVID-19 with mild or no symptoms at all, IgG antibodies could only be detected after 6 weeks (data not shown). In terms of specificity, cross-reacting antibodies of endemic coronavirus infected individuals or of individuals with other active infectious diseases (e.g. EBV or CMV) are a known phenomenon (9). The examined assays in our study demonstrated a good specificity. Only the Vircell ELISA generated one positive result for one HCoV-229E sample, whereas the Euroimmun ELISA generated only one borderline result for the HCoV-OC43 sample and the IFA an unspecific signal in one EBV sample. For the Assure Tech Rapid Test, no cross-reactions were observed, however, a larger sample size would be needed to get a clearer picture. The cross-reactivity of the SARS-CoV samples from the outbreak of 2003 in the Vircell ELISA and IFA are of less importance as the virus is known to be eradicated. Nonetheless, as a false positive result might give a false sense of security, efforts should be made to further improve the specificity of the available assays. All in our study examined assays are eligible for the detection of SARS-CoV-2 IgG antibodies, indicating a potential protective immunity. Ideally, to get the maximum sensitivity, testing should be performed in the later phase of infection (≥ 10 days) after PCR-confirmation or disease onset of COVID-19. The Vircell ELISA, IFA and PRNT demonstrated the highest sensitivity throughout our study. At the moment, however, the PRNT is still the method of choice for questions regarding potential SARS-CoV-2 immunity and should be performed when available.

## Supplementary Material

**TABLE S1.** For sensitivity tested individual follow-up samples of SARS-CoV-2 PCR-confirmed individuals at different time points and generated results.

| Sample Nr. | Day after confirmed SARS-CoV-2 PCR | Euroimmun (ELISA) S/CO | Vircell (ELISA) S/CO | IFA (in-house) qual. | Assure Tech (Rapid Test) qual. | PRNT Titer |
| --- | --- | --- | --- | --- | --- | --- |
| 1 | 5 | <0.8 | 1.0 | neg. | neg. | neg. |
| 2 | 6 | 6,2 | 3.6 | pos. | pos. | 1:160 |
| 3 | 6 | <0.8 | <0.4 | neg. | neg. | neg. |
| 4 | 6 | <0.8 | 1.1 | pos. | pos. | 1:40 |
| 5 | 6 | 11 | 3.3 | pos. | pos. | 1:1280 |
| 6 | 6 | 10.1 | 4.1 | pos. | pos. | 1:640 |
| 7 | 7 | 15.8 | 4.2 | pos. | pos. | 1:640 |
| 8 | 7 | 12.3 | 5.1 | pos. | pos. | 1:1280 |
| 9 | 7 | 14.1 | 5.1 | pos. | pos. | 1:1280 |
| 10 | 8 | <0.8 | <0.4 | neg. | neg. | neg. |
| 11 | 8 | <0.8 | <0.4 | neg. | neg. | neg. |
| 12 | 8 | <0.8 | <0.4 | pos. | neg. | neg. |
| 13 | 8 | 10 | 5 | pos. | pos. | 1:640 |
| 14 | 8 | 14.5 | 5.1 | pos. | pos. | 1:640 |
| 15 | 8 | 13.8 | 5.1 | pos. | - | 1:320 |
| 16 | 9 | <0.8 | <0.4 | pos. | neg. | 1:10 |
| 17 | 9 | 15.7 | 4.9 | pos. | pos. | 1:1280 |
| 18 | 10 | 1.13 | 2.7 | pos. | pos. | 1:80 |
| 19 | 10 | neg | 1.3 | pos. | neg. | 1:80 |
| 20 | 10 | 5.2 | 2.5 | pos. | pos. | 1:320 |
| 21 | 10 | 6.4 | 1.2 | pos. | pos. | 1:640 |
| 22 | 10 | 16.2 | 4.4 | pos. | pos. | 1:1280 |
| 23 | 11 | 17 | 5.1 | pos. | pos. | 1:640 |
| 24 | 13 | 16.6 | 5.1 | pos. | pos. | 1:1280 |
| 25 | 13 | 15.6 | 4.4 | pos. | pos. | 1:160 |
| 26 | 14 | 6.14 | 4.6 | pos. | pos. | 1:320 |
| 27 | 14 | 17 | 5.1 | pos. | pos. | 1:1280 |
| 28 | 16 | 7.2 | 4.8 | pos. | pos. | 1:640 |
| 29 | 16 | 16.2 | 5.1 | pos. | pos. | 1:640 |
| 30 | 16 | 14.8 | 5.1 | pos. | pos. | 1:640 |
| 31 | 17 | 14.8 | 3.7 | pos. | pos. | 1:640 |
| 32 | 17 | 13.4 | 3.9 | pos. | pos. | 1:1280 |
| 33 | 18 | 13 | 5.1 | pos. | pos. | 1:1280 |

Euroimmun (S/CO <0.8 = negative, 0.8-<1.1 = equivocal, ≥ 1.1 = positive), Vircell (S/CO <0.4 = neg., 0.4-0.6 = equivocal, >0.6 = pos.); pos., positive; neg., negative; -, not tested.

**TABLE S2.** For specificity tested follow-up samples of individuals with selected PCR- or serologically-confirmed infections and generated results.

| Sample Nr. | Recently PCR-/serologically-confirmed infected with | Euroimmun (ELISA) S/CO | Viracell (ELISA) S/CO | IFA (in-house) qual. | Assure Tech (Rapid Test) qual. |
| --- | --- | --- | --- | --- | --- |
| 1 | HCoV-OC43 | neg. | neg. | neg. | neg. |
| 2 | HCoV-OC43 | 0.9 | neg. | neg. | neg. |
| 3 | HCoV-OC43 | neg. | neg. | neg. | neg. |
| 4 | HCoV-OC43 | neg. | neg. | neg. | neg. |
| 5 | HKU 1 | neg. | neg. | neg. | neg. |
| 6 | SARS-CoV-1 | neg. | 2.2 | pos. | - |
| 7 | SARS-CoV-1 | neg. | 3.8 | pos. | - |
| 8 | SARS-CoV-1 | neg. | 3.9 | pos. | - |
| 9 | SARS-CoV-2 neg. | neg. | neg. | neg. | - |
| 10 | SARS-CoV-2 neg. | neg. | neg. | neg. | - |
| 11 | SARS-CoV-2 (neg.) | neg. | neg. | - | - |
| 12 | SARS-CoV-2 + Multiplex* neg. | neg. | neg. | neg. | - |
| 13 | HCoV-229E | neg. | neg. | neg. | - |
| 14 | HCoV-229E | neg. | 1.5 | neg. | - |
| 15 | HCoV 229E + Parainfluenza Virus Type 3 | neg. | neg. | neg. | neg. |
| 16 | HCoV-229E | neg. | neg. | neg. | neg. |
| 17 | HCoV-229E | neg. | neg. | neg. | neg. |
| 18 | HCoV-NL63 + Enterovirus/Rhinovirus | neg. | neg. | neg. | neg. |
| 19 | HCoV-NL63 | neg. | neg. | - | - |
| 20 | CMV (+ IgM antibody pos.) | neg. | neg. | neg. | neg. |
| 21 | CMV (+IgM antibody pos.) | neg. | neg. | neg. | neg. |
| 22 | CMV (+ IgM antibody pos.) | neg. | neg. | - | - |
| 23 | EBV-VCA-IgM pos. | neg. | neg. | neg. | neg. |
| 24 | EBV (+ -VCA-IgM antibody pos.) | neg. | neg. | neg. | neg. |
| 25 | EBV-VCA-IgM antibody pos . | neg. | - | neg. | - |
| 26 | EBV-VCA-IgM antibody pos . | neg. | - | unsp. | - |

Euroimmun (S/CO <0.8 = negative, 0.8-1.1 = equivocal, ≥ 1.1 = positive); Viracell (S/CO <0.4 = neg., 0.4-0.6 = equivocal, >0.6 = pos.); pos., positive; neg., negative; unsp., unspecific; \*Biofire® Filmarray® 20 Target Respiratory Panel (bioMérieux, Nürtingen, Baden-Württemberg, Germany); -, not tested.

